# A new prognostic model for pancreatic cancer based on pyroptosis-related genes

**DOI:** 10.1101/2021.11.08.467376

**Authors:** Liukai Ma, Mengyao Wang, Ruoling Jia, Cheng Yan

## Abstract

Pancreatic cancer is one of the most common malignant tumors of the digestive tract. It is known as the “king of cancer” in the field of cancer, and is one of the worst prognosis malignant tumors. Pyroptosis is a kind of programmed cell death, which can promote the inflammatory response of cells. Studies have shown that the effect of pyroptosis-related genes in cancer is significant. However, the role of pyroptosis in pancreatic cancer is not clear. The aim of this study is to establish a prognostic model based on pyroptosis. The gene expression and clinical data of pancreatic cancer patients were obtained from TCGA and verified in GEO. The differential expression of 33 pyroptosis-related genes in pancreatic cancer and normal tissues was analyzed, of which 6 genes were up-regulated and 12 genes were down regulated. Then, it was analyzed that pyroptosis-related genes were mainly enriched in the defense against bacteria and pyroptosis pathways. A concise and reliable model is established by lasso-cox regression analysis. Km curve shows that there are differences between high-risk group and low-risk group. And the nomogram has reliable prediction ability. In conclusion, pyroptosis an important role in pancreatic cancer, which can be used for the prediction of pancreatic cancer and provide a new perspective for the treatment of pancreatic cancer.

**AUTHOR APPROVALS:** The authors have seen and approved the manuscript, and that it hasn’t been accepted or published elsewhere.

**COMPETING INTERESTS:** The authors declare no competing interests.

## INTRODUCTION

Pancreatic cancer remains one of the deadliest malignancies, recording 432,242 new deaths in 2018, with 458,918 new pancreatic cancer cases reported globally[1]. Pancreatic cancer incidence and mortality rate are nearly equivalent and the 5-year survival rate is estimated at only 1%[2]. Most patients already have advanced disease at the time of diagnosis. Currently, treatment of pancreatic cancer is dependent on surgical resection. However, up to 80% of pancreatic cancers are unresectable for their highly malignant and early metastasis[3]. Even after resection of pancreatic cancer, the prognosis of pancreatic cancer is poor[4]. Therefore, it is important to establish a good prognostic model.

Pyroptosis, also known as cellular inflammatory necrosis, is a novel form of programmed cell death[5]. Studies have shown that the serine protease granzyme A released by lymphocytes such as cytotoxic T lymphocytes (CTLs) and NK cells can enter tumor cells through perforin on the surface of tumor cells, specifically and efficiently cut GSDMB protein, resulting in tumor cell scorch[6]. Caspase-1–deficient mice demonstration increased colonic epithelial cell proliferation in early stages of injury-induced tumor formation and reduced apoptosis in advanced tumors[7]. These studies have proved that pyroptosis plays a dual role in tumors.

To further improve the effectiveness of pancreatic cancer treatment and formulate accurate treatment strategies, oncologists need to identify the prognosis of patients with pancreatic cancer. At present, there is no prognostic model of pyroptosis gene in pancreatic cancer. Therefore, exploring the related molecular biomarkers of pyroptosis would have an attractive value in estimating and treating pancreatic cancer, which may be an important therapeutic trend of pancreatic cancer.

## METHODS

### Data acquisition and pretreatment of pancreatic cancer

We downloaded the RNA sequencing data and clinical data of 178 pancreatic cancer patients from TCGA in August 19, 2021. In addition, two sets of pancreatic cancer RNA-Seq data and corresponding clinical information were downloaded in GEO as external validation cohort (https://www.ncbi.nlm.nih.gov/geo/query/acc.cgi, ID:GSE71729; https://www.ncbi.nlm.nih.gov/geo/query/acc.cgi,ID:GSE57495). Two sets of data in GEO are extracted, annotated and normalized by ((https://www.omicstudio.cn/tool)). The gene expression matrix after the combination of GSE71729 and GSE57495 was obtained.

### Identification of Differential Expression pyroptosis-related genes and Enrichment Analysis

We analyzed differential genes in 178 patients in the TCGA cohort and 167 normal tissues in the GTEX cohort using the deseq2 package. The thresholds were adjusted P < 0.05 and | logFC | > 1.2. The volcano map and heat map of differentially expressed genes were drawn by LiancHuan cloud tool, and the protein-protein interaction (PPI) was drawn by string database (https://string-db.org/), the box diagram is drawn by package R (ggpubr). The differentially expressed genes were enriched by GO and KEGG by R software. The R packages used include go: goplot, KEGG: clusterprofiler, org.hs.eg.db, enrich plot and ggplot2. In addition, we also did GASE enrichment analysis.

### Construction of pyroptosis-related genes risk score and prognosis model

We first made a single factor COX regression analysis of genes related to focal burn. The significant filtering standard was pFilter=0.05 (R package: survival). After preliminary screening, we excluded genes associated with pancreatic cancer prognosis. After that, the prognosis model was constructed by lasso regression analysis, and through scoring formula 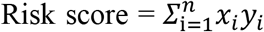 (x is the coef value obtained by lasso regression analysis and Y is the gene expression), and the risk score was given to the patients. According to the median of risk, the patients in the TCGA cohort were divided into high risk group and low risk group. Similarly, the patients in the GEO cohort were divided into high-risk group and low risk group according to the median TCGA queue risk. The “survival” package of R software was used for Kaplan–Meier survival analysis. The predictive accuracy of each gene and the risk score were evaluated by performing time receiver-operating characteristic (ROC) analysis. We extracted the clinical information (age, grade, rade, stage, T and N) of patients in the TCGA cohort .These variables were analysed in combination with the risk score in our regression model. Univariate and multivariable Cox regression models were employed for the analysis.

### The Construction of Nomogram and Calibration Curves

A clinical practical nomogram was established to predict individual survival probability by the “rms” package of R software. To assess the consistency between actual and predicted survival in the nomogram, calibration curves for predicting 1-, and 3-year survival rate were also drawn. The prediction ability of signature was assessed by area under ROC curve.

## RESULTS

### Screening and identification of differentially expressed genes related to PAAD

We obtained 178 PAAD patients from TCGA and 167 normal tissues from GTEX. The differential expression of 33 pyroptosis-related genes was analyzed by r-packet (deseq2). It was found that 18 pyroptosis-related genes were differentially expressed (P < 0.05). Obtain volcano map (| log2 (fold change) | > 1.2, FDR p-value < 0.05) on LianChuan biological cloud platform (Figure.1A). Six genes were up-regulated and 12 genes were down regulate. The RNA expression levels of these genes are shown in the figure. To further explore the interactions of these pyroptosis-related genes, we conducted a protein–protein interaction (PPI) analysis, and the results are shown. The minimum required interaction score for the PPI analysis was set at 0.4 (medium confidence) (Figure.1B). At the same time, the heat map also proves that 6 genes are adjusted, and 12 genes are lowered (Figure.1C). At the same time, we also made a mutation of the focal pyroptosis-related genes related to PAAD. The results showed that the expression of the pyroptosis-related genes was more valuable than the mutation (Figure.1E).

**Figure.1.**
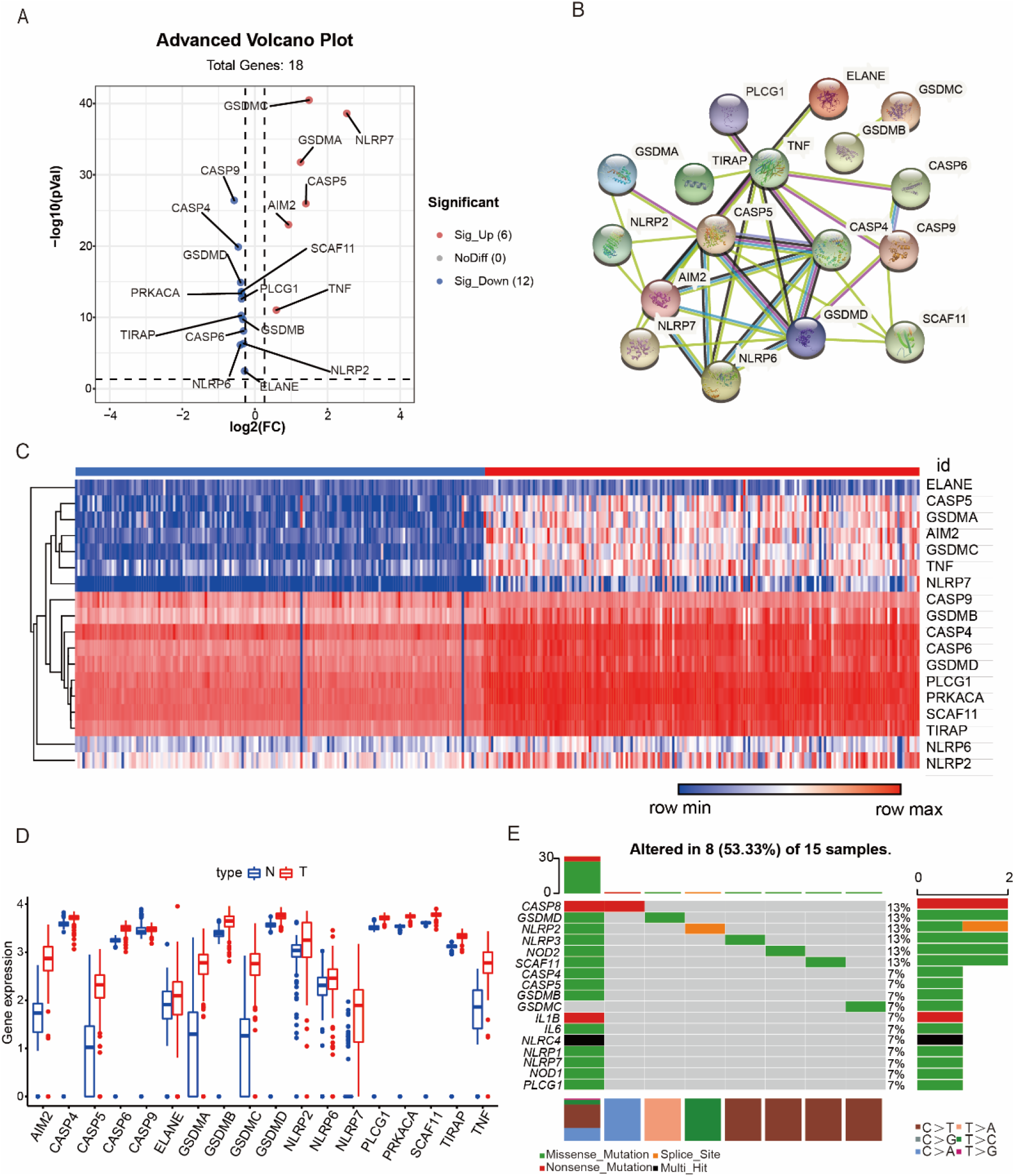
The differentially expressed pyroptosis-related genes between lung adenocarcinoma and normal tissues. (A) Volcanic map representation differentially expressed pyroptosis-related genes, with red dots indicating high expression and blue dots indicating low expression. (B) Protein-protein interactions (PPI) network showing the interactions of the pyroptosis-related genes (interaction score=0.4). (C) Heatmap of differentially expressed pyroptosis-related genes, with red indicating high expression and green indicating low expression. (D) Box plot of DEAGs, with the red boxes representing the tumor group and blue boxes representing the normal group. Differential expression of the 18 selected genes between normal and pancreatic cancer tissues. (E) Differential somatic mutation analysis between high-risk group and low-risk group in The Cancer Genome Atlas (TCGA) cohort.

### GO and KEGG pathway analysis

In order to further understand the function of genes related to pyroptosis, we analyzed these genes by go and KEGG. We found that these 33 cases were mainly related to the formal regulation of cytokine production and the defense response to bacteria (Figure.2A). In addition, KEGG pathway analysis showed that pyroptosis-related genes were involved in platinum drug resistance, apoptosis-multiple species, ErbB signaling pathway and apoptosis (Figure.2B)

**Figure.2.**
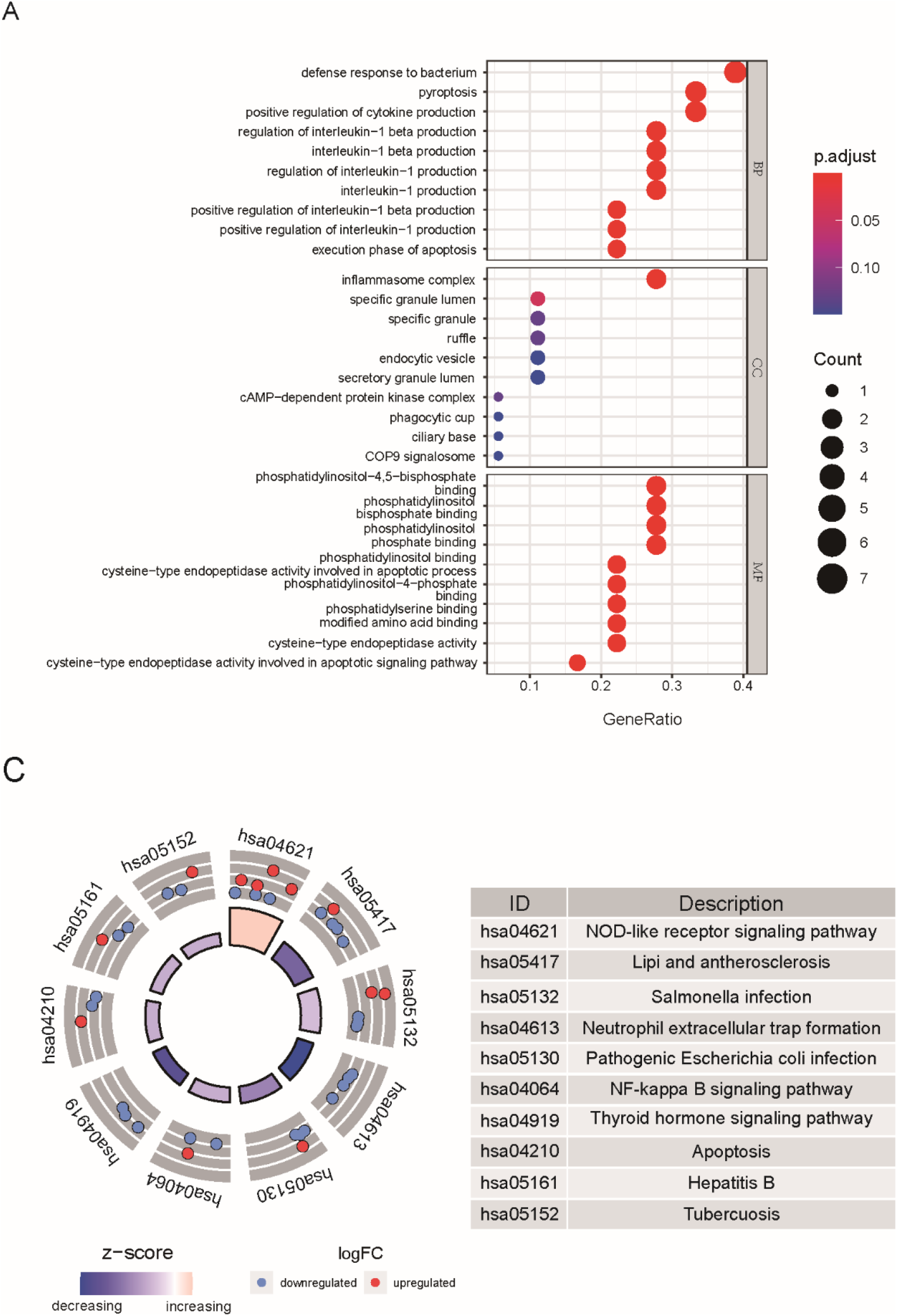
Functional enrichment analyses of gene ontology (GO) and Kyoto Encyclopedia of Genes and Genomes (KEGG). (A) Bubble graph for GO enrichment (the bigger bubble means the more genes enriched, and the increasing depth of red means the differences were more obvious; q-value: the adjusted p-value). (B) The circular scatter plot of enriched KEGG terms.

### Construction of differential gene prediction model for pancreatic cancer

A total of 170 patients with pancreatic cancer who had survival data were analyzed by univariate regression analysis. We screened 9 genes related to blepharoptosis based on P value less than 0.05 (IL18, GSDMC, NLRP2, PLCG1, GPX4, CASP8, PRKACA, NLRP1 and CASP4). Among them, and higher expressions of IL18, GSDMC, NLRP2, CASP8, CASP4 were associated with increased risk (HR > 1), while upregulated expressions of PLCG1, GPX4, PRKACA, NLRP1 were correlated with lower risk (HR < 1) ((Figure.3A). Subsequently, LASSO Cox regression analysis retrieved 5 genes for prognostic model construction based on the optimum λ value (Figure.3C, 3D).

**Figure.3.** Forest map of heat sag related genes in patients with pancreatic cancer. And, construction of pyroptosis-related gene risk signature in TCGA cohort. (A) Univariate Cox regression analysis. (C) Cross-validation for tuning the parameter selection in the LASSO regression. (D) LASSO regression of the 9 OS-related genes.

### Establishment of prognostic model in TCGA cohort

In order to better construct the prognostic model related to pyroptosis genes. We calculated the risk score for each patient according to the following formula and divide 170 patients into high-risk group(n = 85) and low-risk group (n = 85) according to the median value of risk score (Figure.4A). Compared with the low-risk group, the high-risk group had more deaths and shorter survival time. And risk of death was significantly increased in the high-risk group (Figure. 4C). IL18, CASP4, GSDMC and NLRP2 were up-regulated in high-risk group and NLRP1 was down regulated in high-risk group (Figure. 4D). The Kaplan–Meier curve demonstrated that patients in the high-risk group have a poorer prognosis (P < 0.05, Figure.4B).

**Figure.4.**
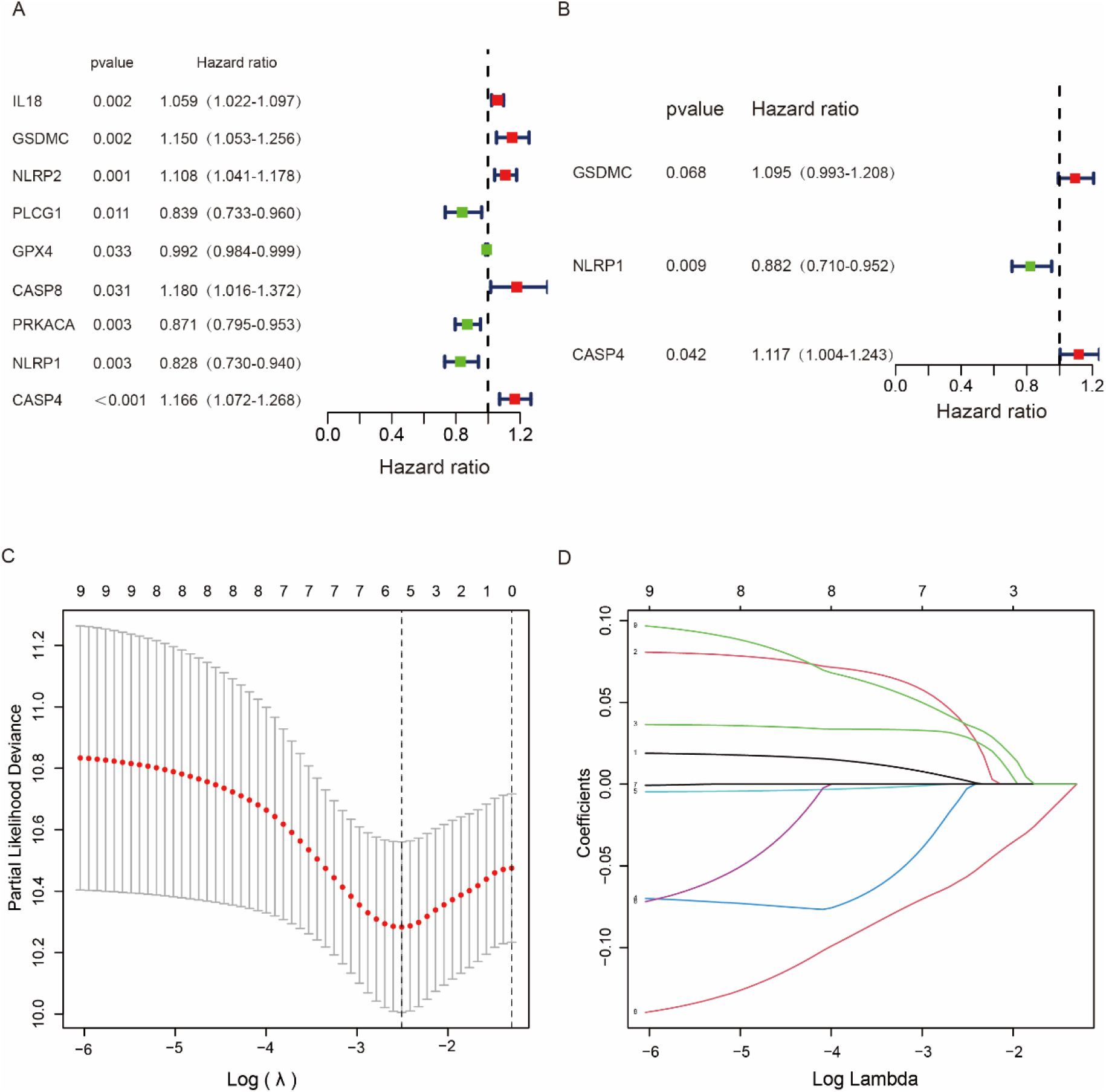
Construction of risk signature in the TCGA cohort. (A) Distribution of patients based on the risk score. (B) Kaplan–Meier curves for the overall survival of patients in the high- and low-risk groups. (C) Survival status of patients with pancreatic cancer. (D) Expression of five pyroptosis-related risk genes.

### External validation of the risk signature

To verify the accuracy of the prognostic model established by TCGA, we obtained 186 pancreatic cancer patients from GEO and made a risk score with the same formula. Based on the median risk score in the TCGA cohort, 99 patients in the GEO cohort were classified into the low-risk group, while the other 87 patients were classified into the high-risk group (Figure. 5A). Patients in the low-risk subgroup (Figure. 5C, on the left side of the dotted line) were found to have longer survival times and lower death rates than those in the high-risk subgroup. IL18, CASP4, GSDMC and NLRP2 were up-regulated in high-risk group and NLRP1 was down regulated in high-risk group (Figure. 5D). In addition, Kaplan–Meier analysis also indicated a significant difference in the survival rate between the low-risk and high-risk groups (P<0.05, Figure. 5B). The results are the same as those in the TCGA cohort.

**Figure.5.**
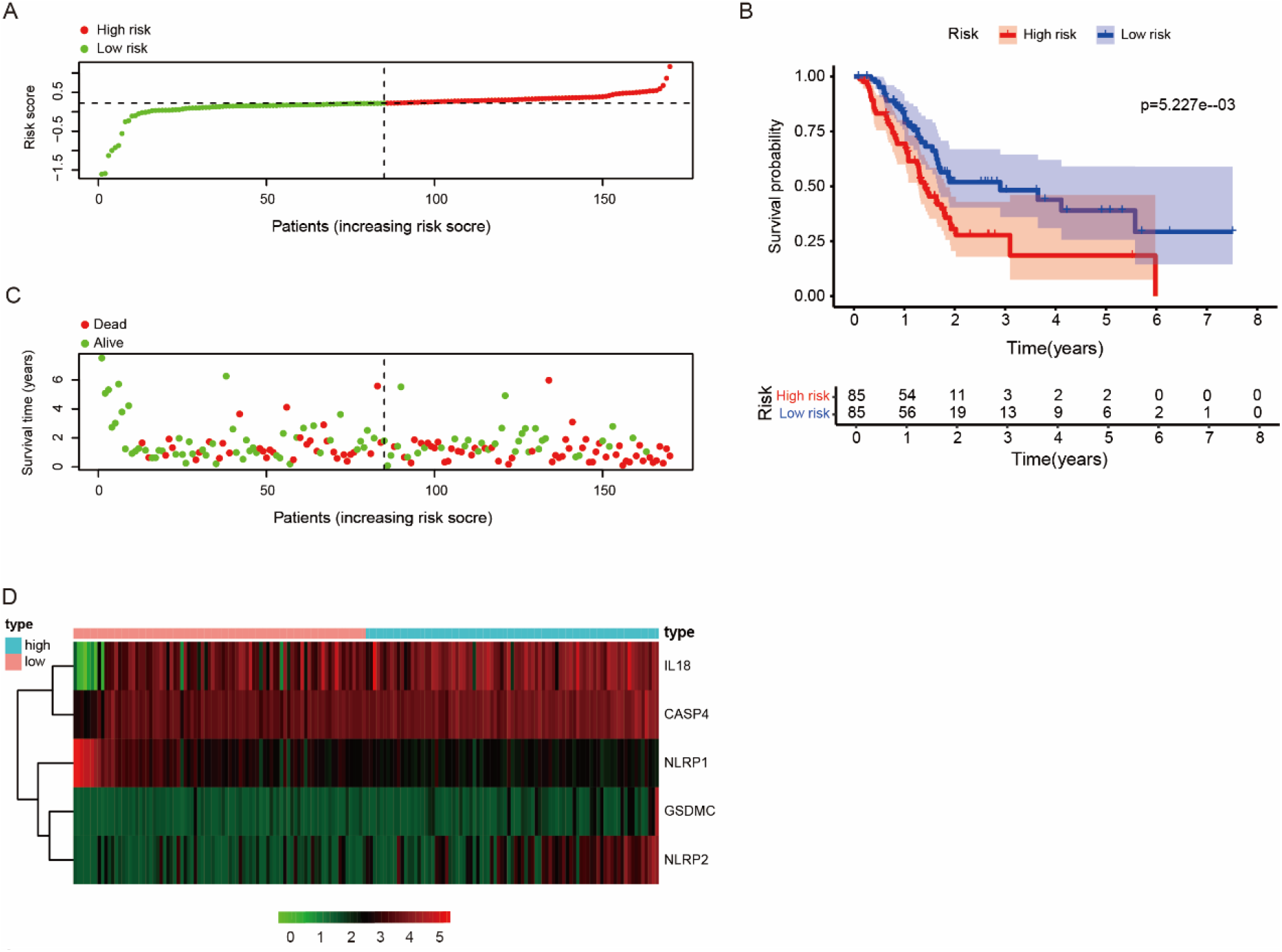
Validation of the risk model in the GEO cohort. (A) Distribution of risk score. (B) Kaplan–Meier curves for the overall survival of patients in the high- and low-risk groups. (C) Survival status of pancreatic cancer patients in GEO cohort. (D) Expression heat map of 5 differential expression of pyroptosis-related genes.

### Independent prognostic analysis of TCGA

In order to verify whether the prognostic model we constructed can be independent of other clinical traits as an independent prognostic factor, univariate and multivariate Cox regression analyses were performed. Univariate Cox results showed that age, N stage and risk score were correlated with survival (P < 0.05), and risk score was significantly correlated with survival status (P < 0.001, Figure. 6A). Multivariate Cox regression analysis showed that the risk score was significantly correlated with overall survival (P < 0.001, Figure. 6B). The results showed that risk score was an independent prognostic index.

**Figure.6.**
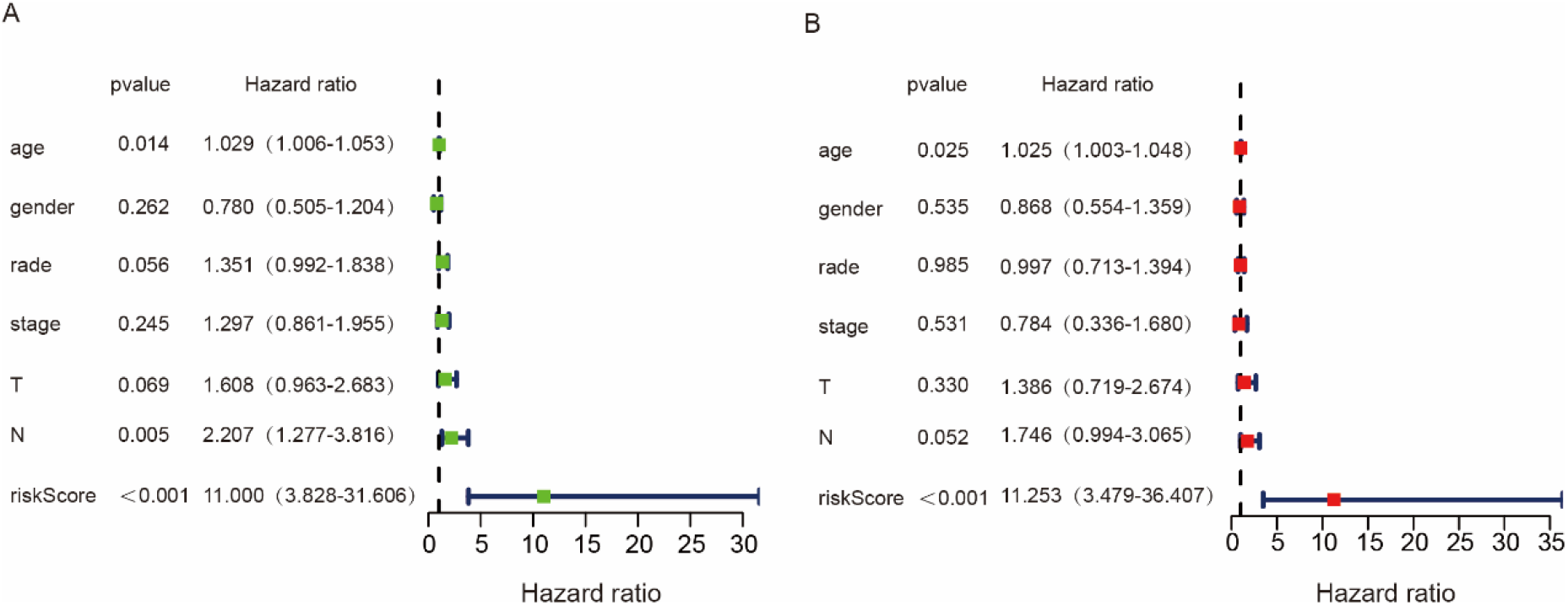
Independent prognostic analysis. (A) Univariate independent prognostic analysis. (B) Multivariate independent prognostic analysis.

### Relationship between pyroptosis-related genes expression and clinical information

Nlrp1 (Figure. 7A, p=0.022) was negatively correlated with grade, N (Figure. 7B, p=0.022) and T stage (Figure. 7C, p=0.046). Nlrp2 was positively correlated with grade (Figure. 7D, p=0.022). GSDMC was negatively correlated with stage (Figure. 7E, p=0.010). CASP4 was positively correlated with grade (Figure. 7F, p=0.012). Risk scores were positively correlated with grade (Figure. 7G, p=5.002e−04).

**Figure.7.**
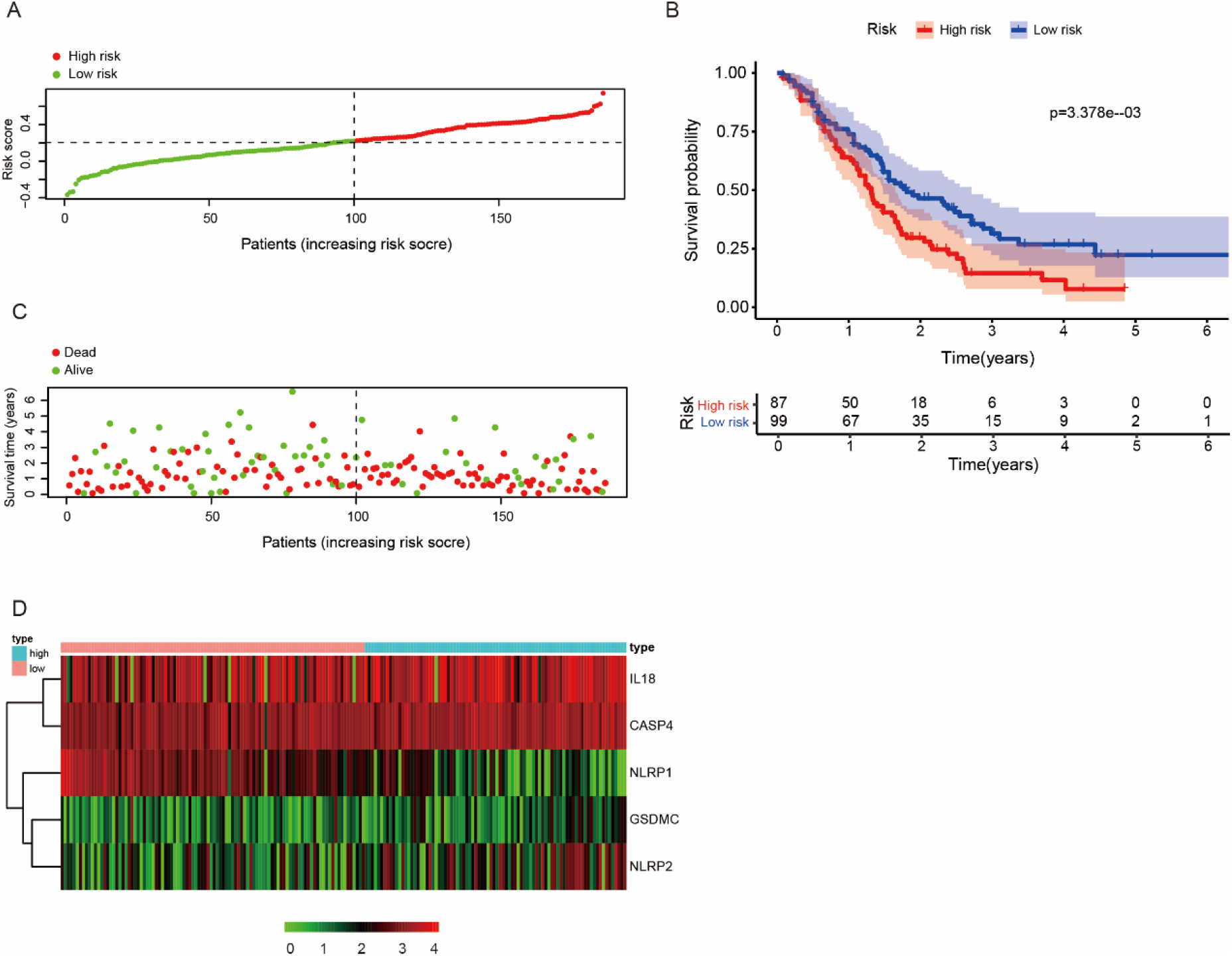
Clinicopathological significance of pyroptosis-related genes and risk score in pancreatic cancer. (A,B,C) Clinicopathological correlation of NLRP1 in pancreatic cancer. (D)Clinicopathological correlation of NLRP2 in pancreatic cancer. (E) Clinicopathological correlation of GSDMC in pancreatic cancer. (F) Clinicopathological correlation of GASP4 in pancreatic cancer. (E) Clinicopathological correlation of risk score in pancreatic cancer.

### Construction and Validation of the Nomogram

In order to predict the prognosis of pancreatic cancer more accurately, we used clinical information and risk scores to make a nomogram. The survival probability of pancreatic cancer patients in 1 and 3 years is evaluated. The nomogram showed that our risk score was the most important factor among the various clinical parameters (Figure. 8A). In order to determine the accuracy of the constructed nomogram, we constructed the calibration curve, and the calibration curve showed that the prediction of the line map was consistent with the actual survival of the patients of pancreatic cancer (Figure. 8B, C). Finally, the predictive abilities of the nomogram were analyzed by the AUC values (AUC of 1-year OS=0.664, Figure 8D).

**Figure.8.**
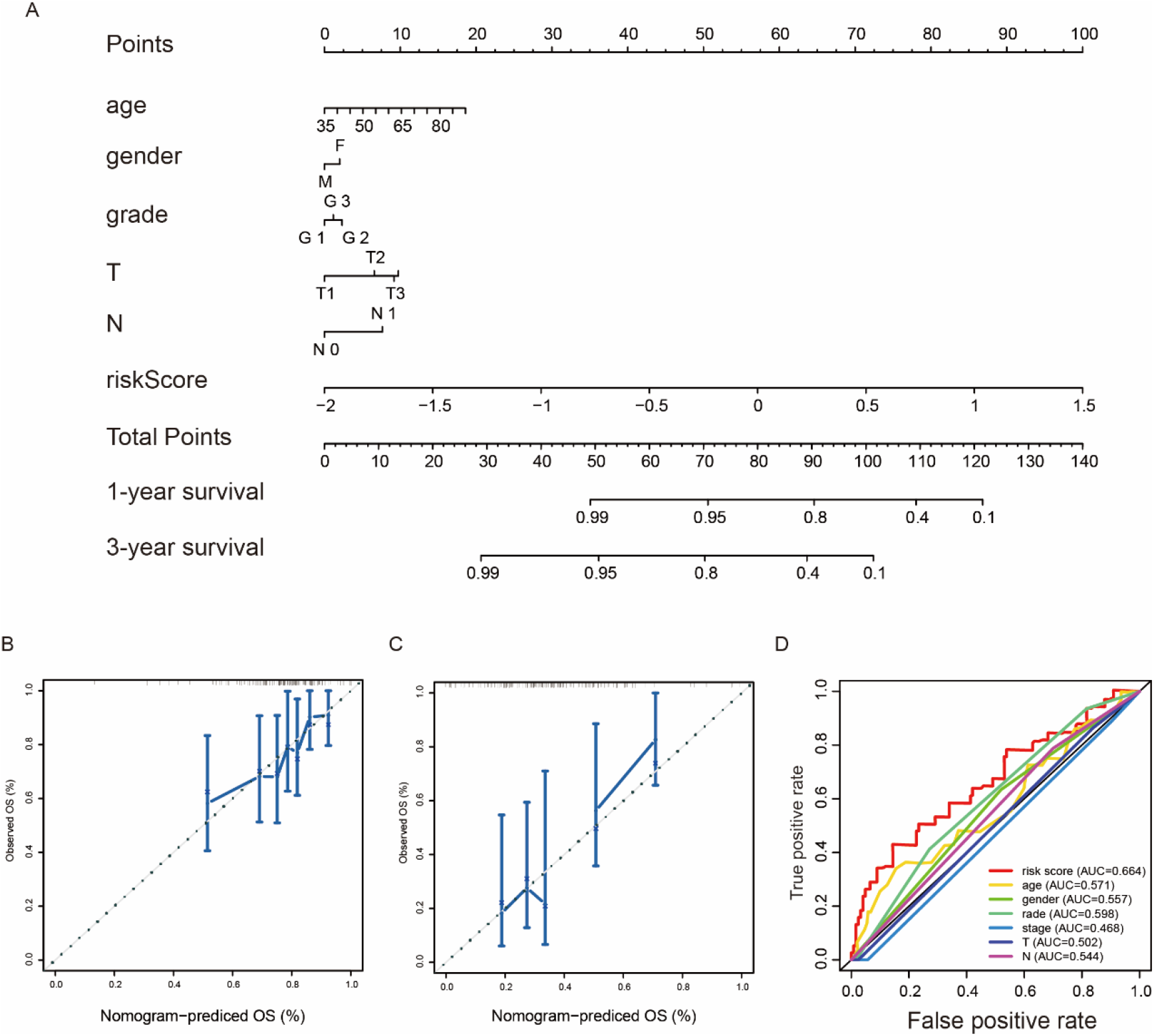
Nomogram to predict the probability of patients with pancreatic cancer. (A) A nomogram integrating the signature risk score with the pathologic characteristics. (B, C) The calibration plots for predicting patient 1-, 3-year OS. (D) The ROC curve of OS for risk score, age, gender, stage, N stage and T stage at 1-year.

### Gene Set Enrichment Analysis

Finally, we performed gene enrichment analysis between high-risk and low-risk groups. The results showed that the main enrichment pathway of high-risk group was apoptosis, tight junction, spliceosome, axon guidance, natural killer cell mediated cytotoxicity. The main enrichment pathways of low risk are neuroactive ligand receptor interaction, taste transduction, autoimmune thyroid disease, chemokine signaling pathway, focal adhesion (Figure 9A).

**Figure.9.**
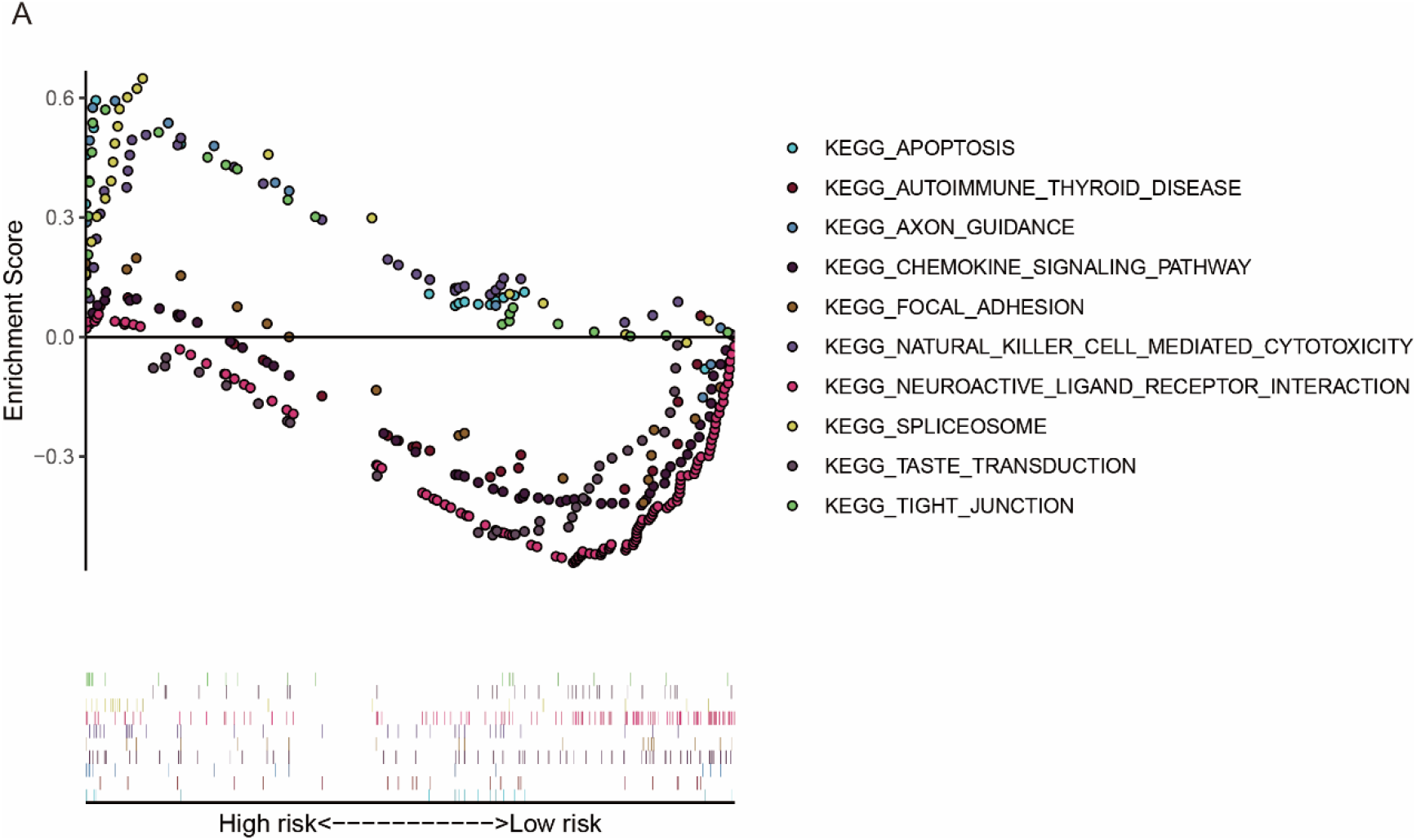
Multi GSEA enrichment map. (A) GSEA analysis of the differentially expressed genes between high- and low-risk groups.

### Single gene Kaplan Meier curve

In addition, we also constructed a single gene model. The results showed that there was a significant difference between NLRP1 and NLRP2 (P < 0.05,Figure 10C,D). There was no significant difference in GSDMC, IL18 and CASP4 (P > 0.05, Figure 10A, B, E).

**Figure.10.**
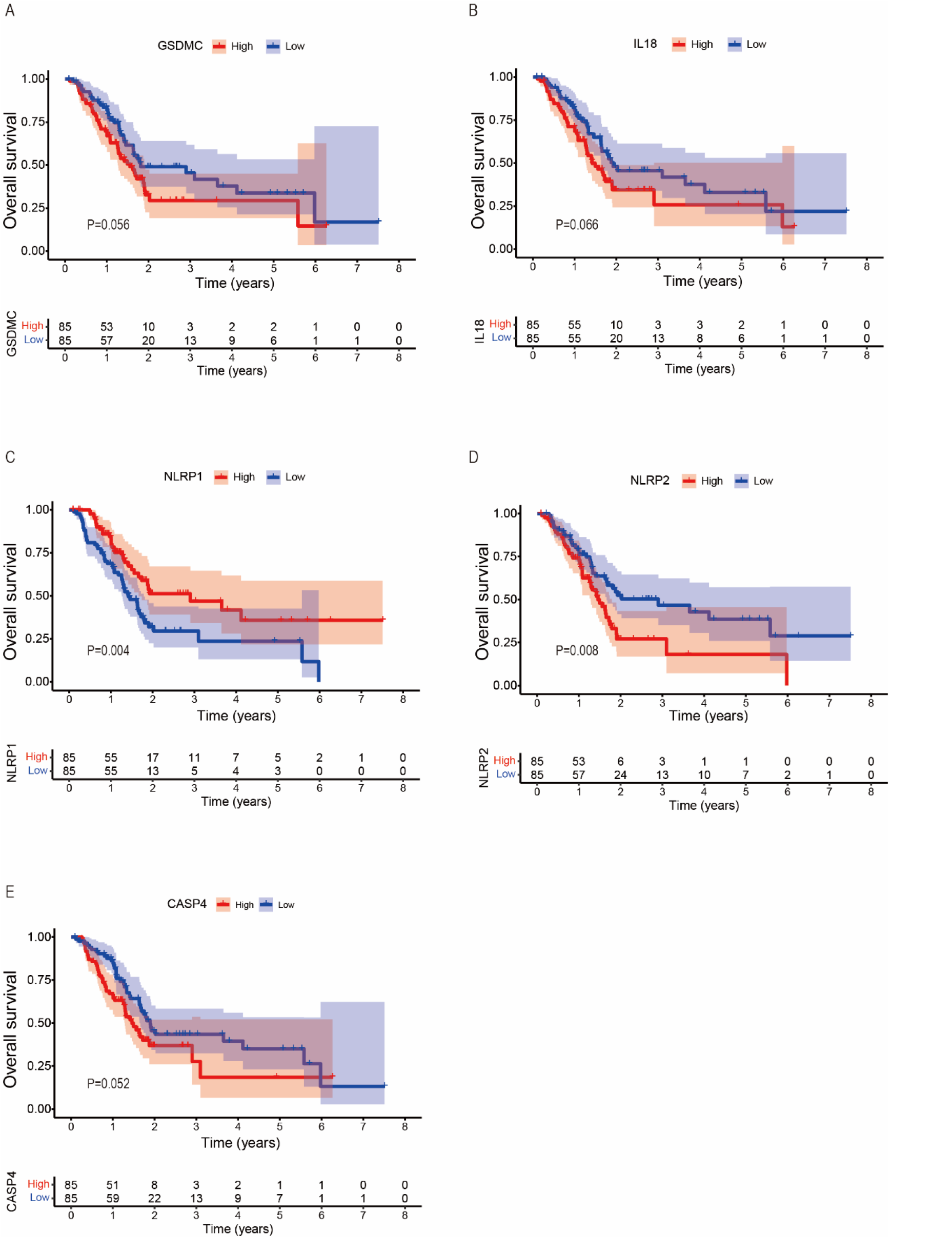
Single gene Kaplan Meier curve. (A,B,C,D,E,) Single gene Kaplan Meier curve of GSDMC, IL18, NLRP 1, NLRP 2 and CASP4.

## DISSCUSION

Pyroptosis is a kind of caspase-1 or caspase-11-dependent programmed cell death[8, 9]. In recent years, there are more and more studies on cell pyroptosis, and the role of pyroptosis in cancer cells is gradually clear. Studies have shown that pyroptosis genes play different roles in different cancers. Cell death releases inflammatory factors, which provides a suitable living environment for tumor cells[10]. Pyroptosis can promote tumour cell death, making pyrolysis a potential prognostic and therapeutic target for cancer[11]. However, the role of pyroptosis gene in pancreatic cancer is not clear. The purpose of this study is to elucidate the role of pyroptosis gene in pancreatic cancer.

In recent years, with the rapid development of cancer biology knowledge, there are many researches on pancreatic cancer metastasis, new biomarkers, and new prognostic markers[12-16]. These provide good prognostic models for pancreatic cancer patients. Compared with these studies, our research mainly demonstrates the effect of differential expression of pyroptosis genes on pancreatic cancer patients. Although there is a certain gap with other prognostic models, the results suggest that the pyroptosis gene has a good prognosis in pancreatic cancer patients.

We constructed a prognostic risk model of 5 genes (IL18, CASP4, NLRP1, NLRP2, GSDMC) through lasso Cox regression analysis, and different risk groups have different survival results. This result has also been verified in the validation group, which proves the effectiveness of the model. The nomogram also well predicted the 1-year and 3-year overall survival rates.

We found that the expression of IL18, CASP4, GSDMC and NLRP2 in high-risk group was higher than that in low-risk group, but the expression of these genes was different in other diseases. For example, the lack of IL18 can lead to seizures in patients with Alzheimer’s disease [17]. High expression of CASP4 will lead to poor prognosis in patients with renal cell carcinoma[18], but low expression of CASP4 will lead to poor prognosis in patients with esophageal squamous cell carcinoma[19]. Overexpression of GSDMC inhibits TGFBr2 (transforming growth factor β receptor type ii) promotes the proliferation of colorectal cancer cells and leads to poor prognosis in patients with lung adenocarcinoma[20, 21]. This shows that the same gene plays different roles in different diseases.

It is worth mentioning that in the study, we found that NLRP1 is one of the important genes. NLRP1 is a key regulator of the innate immune system. Patients with vitiligo increase interleukin-1 through NLRP 1 inflammatory corpuscles β the high expression of[22]. NLRP1 can aggravate inflammatory bowel disease by limiting the production of butyric acid[23]. The high expression of NLRP1 can cause psoriasis [24]. In addition, NLRP1 is also closely related to type 1 diabetes [25]. These studies indicate that high expression of NLRP1 aggravates the disease, but in our study, high expression of NLRP1 is more beneficial to patients. This shows that NLRP1 plays different roles in different cancers. Although NLRP1 has been widely studied, the role of NLRP1 in pancreatic cancer is limited. In our research, NLRP1 is one of the biomarkers of prognosis in pancreatic cancer.

We analyzed the effect of single gene on the survival rate of pancreatic cancer patients. We found that NLRP1 gene had a significant effect on pancreatic cancer patients. Moreover, the prognosis of pancreatic cancer is better when NLRP1 is overexpressed. This is consistent with the results of figure4D and figure5D, indicating that the role of NLRP1 in pancreatic cancer deserves further study.

However, our study has potential limitations. First, we do not have enough clinical data to verify the expression of M stage in the prognostic model. Secondly, the model we established needs to be further verified by in vitro and in vivo experiments to increase the reliability of the prognostic model.

In conclusion, we constructed a prognostic risk model for pancreatic cancer using 5 differentially expressed pyroptosis genes, which can accurately predict the prognosis of patients with pancreatic cancer. It is demonstrated that pyroptosis gene is one of the biomarkers of survival rate in pancreatic cancer patients.

## REFERENCES

1. Bray F, Ferlay J, Soerjomataram I, Siegel RL, Torre LA, Jemal A: Global cancer statistics 2018: GLOBOCAN estimates of incidence and mortality worldwide for 36 cancers in 185 countries. CA Cancer J Clin 2018, 68(6):394–424.

2. Tamburrino A, Piro G, Carbone C, Tortora G, Melisi D: Mechanisms of resistance to chemotherapeutic and anti-angiogenic drugs as novel targets for pancreatic cancer therapy. Front Pharmacol 2013, 4:56.

3. Vincent A, Herman J, Schulick R, Hruban RH, Goggins M: Pancreatic cancer. The Lancet 2011, 378(9791):607–620.

4. Seiler C, Gillen S, Schuster T, Meyer zum Büschenfelde C, Friess H, Kleeff J: Preoperative/Neoadjuvant Therapy in Pancreatic Cancer: A Systematic Review and Meta-analysis of Response and Resection Percentages. PLoS Medicine 2010, 7(4).

5. Kovacs SB, Miao EA: Gasdermins: Effectors of Pyroptosis. Trends in Cell Biology 2017, 27(9):673–684.

6. Zhou Z, He H, Wang K, Shi X, Wang Y, Su Y, Wang Y, Li D, Liu W, Zhang Y et al: Granzyme A from cytotoxic lymphocytes cleaves GSDMB to trigger pyroptosis in target cells. Science 2020, 368(6494).

7. Hu B, Elinav E, Huber S, Booth CJ, Strowig T, Jin C, Eisenbarth SC, Flavell RA: Inflammation-induced tumorigenesis in the colon is regulated by caspase-1 and NLRC4. Proceedings of the National Academy of Sciences 2010, 107(50):21635–21640.

8. Chen X, He WT, Hu L, Li J, Fang Y, Wang X, Xu X, Wang Z, Huang K, Han J: Pyroptosis is driven by non-selective gasdermin-D pore and its morphology is different from MLKL channel-mediated necroptosis. Cell Res 2016, 26(9):1007–1020.

9. Fink SL, Cookson BT: Apoptosis, pyroptosis, and necrosis: mechanistic description of dead and dying eukaryotic cells. Infect Immun 2005, 73(4):1907–1916.

10. Huang Y, Zhang Q, Lubas M, Yuan Y, Yalcin F, Efe IE, Xia P, Motta E, Buonfiglioli A, Lehnardt S et al: Synergistic Toll-like Receptor 3/9 Signaling Affects Properties and Impairs Glioma-Promoting Activity of Microglia. J Neurosci 2020, 40(33):6428–6443.

11. Ruan J, Wang S, Wang J: Mechanism and regulation of pyroptosis-mediated in cancer cell death. Chem Biol Interact 2020, 323:109052.

12. Ge JN, Yan D, Ge CL, Wei MJ: LncRNA C9orf139 can regulate the growth of pancreatic cancer by mediating the miR-663a/Sox12 axis. World J Gastrointest Oncol 2020, 12(11):1272–1287.

13. Higuchi T, Yokobori T, Naito T, Kakinuma C, Hagiwara S, Nishiyama M, Asao T: Investigation into metastatic processes and the therapeutic effects of gemcitabine on human pancreatic cancer using an orthotopic SUIT-2 pancreatic cancer mouse model. Oncol Lett 2018, 15(3):3091–3099.

14. Kong L, Liu P, Fei X, Wu T, Wang Z, Zhang B, Li J, Tan X: A Prognostic Prediction Model Developed Based on Four CpG Sites and Weighted Correlation Network Analysis Identified DNAJB1 as a Novel Biomarker for Pancreatic Cancer. Front Oncol 2020, 10:1716.

15. Ma Q, Wu X, Wu J, Liang Z, Liu T: SERP1 is a novel marker of poor prognosis in pancreatic ductal adenocarcinoma patients via anti-apoptosis and regulating SRPRB/NF-kappaB axis. Int J Oncol 2017, 51(4):1104–1114.

16. Wang H, Wang X, Xu L, Lin Y, Zhang J, Cao H: Identification of genomic alterations and associated transcriptomic profiling reveal the prognostic significance of MMP14 and PKM2 in patients with pancreatic cancer. Aging (Albany NY) 2020, 12(18):18676–18692.

17. Tzeng TC, Hasegawa Y, Iguchi R, Cheung A, Caffrey DR, Thatcher EJ, Mao W, Germain G, Tamburro ND, Okabe S et al: Inflammasome-derived cytokine IL18 suppresses amyloid-induced seizures in Alzheimer-prone mice. Proc Natl Acad Sci U S A 2018, 115(36):9002–9007.

18. Meng L, Tian Z, Long X, Diao T, Hu M, Wang M, Zhang W, Zhang Y, Wang J, He Y: Prognostic autophagy model based on CASP4 and BIRC5 expression in patients with renal cancer: independent datasets-based study. Am J Transl Res 2020, 12(11):7475–7489.

19. Shibamoto M, Hirata H, Eguchi H, Sawada G, Sakai N, Kajiyama Y, Mimori K: The loss of CASP4 expression is associated with poor prognosis in esophageal squamous cell carcinoma. Oncol Lett 2017, 13(3):1761–1766.

20. Miguchi M, Hinoi T, Shimomura M, Adachi T, Saito Y, Niitsu H, Kochi M, Sada H, Sotomaru Y, Ikenoue T et al: Gasdermin C Is Upregulated by Inactivation of Transforming Growth Factor beta Receptor Type II in the Presence of Mutated Apc, Promoting Colorectal Cancer Proliferation. PLoS One 2016, 11(11):e0166422.

21. Wei J, Xu Z, Chen X, Wang X, Zeng S, Qian L, Yang X, Ou C, Lin W, Gong Z et al: Overexpression of GSDMC is a prognostic factor for predicting a poor outcome in lung adenocarcinoma. Mol Med Rep 2020, 21(1):360–370.

22. Levandowski CB, Mailloux CM, Ferrara TM, Gowan K, Ben S, Jin Y, McFann KK, Holland PJ, Fain PR, Dinarello CA et al: NLRP1 haplotypes associated with vitiligo and autoimmunity increase interleukin-1beta processing via the NLRP1 inflammasome. Proc Natl Acad Sci U S A 2013, 110(8):2952–2956.

23. Tye H, Yu CH, Simms LA, de Zoete MR, Kim ML, Zakrzewski M, Penington JS, Harapas CR, Souza-Fonseca-Guimaraes F, Wockner LF et al: NLRP1 restricts butyrate producing commensals to exacerbate inflammatory bowel disease. Nat Commun 2018, 9(1):3728.

24. Ciazynska M, Olejniczak-Staruch I, Sobolewska-Sztychny D, Narbutt J, Skibinska M, Lesiak A: The Role of NLRP1, NLRP3, and AIM2 Inflammasomes in Psoriasis: Review. Int J Mol Sci 2021, 22(11).

25. Sun X, Xia Y, Liu Y, Wang Y, Luo S, Lin J, Huang G, Li X, Xie Z, Zhou Z: Polymorphisms in NLRP1 Gene Are Associated with Type 1 Diabetes. J Diabetes Res 2019, 2019:7405120.

